# Megakaryocyte infection by SARS-CoV-2 drives the formation of pathogenic afucosylated IgG antibodies in mice

**DOI:** 10.1101/2023.07.14.549113

**Authors:** Meryem Mabrouk, Farid Jalali, Nabil Zaid, Abdallah Naya, Meriem Khyatti, Loubna Khalki, Fadila Guessous, Younes Zaid

## Abstract

More than 90% of total human plasma immunoglobulin G (IgG) is found in a fucosylated form, but specific IgGs with low core fucosylation (afucosylated IgGs) are found in response to infections with enveloped viruses and to alloantigens on blood cells. Afucosylated IgGs mediate immunopathology in severe COVID-19 and dengue fever in humans. In COVID-19, the early formation of non-neutralizing afucosylated IgG against the spike protein predicts and directly mediates disease progression to severe form. IgG lacking core fucosylation causes dramatically increased antibody-dependent cellular toxicity mediated by intense FcγR-mediated stimulation of macrophages, monocytes, natural killer cells, and platelets. The mechanism and the context within which afucosylated IgG formation occurs in response to enveloped virus antigens have remained elusive thus far in COVID-19, dengue fever, and other infections. This study demonstrates that administration of human bone marrow megakaryocytes infected by SARS-CoV-2 into the circulation of K18-hACE2 transgenic mice drives the formation of pathogenic afucosylated anti-spike IgG antibodies, and is sufficient to reproduce severe COVID-19 manifestations of pulmonary vascular thrombosis, acute lung injury, and death in mice.

## Introduction

The coronavirus SARS-CoV-2 is the causative agent of the global coronavirus 2019 (COVID-19) pandemic which has caused a severe threat to public health since its inception. SARS-CoV-2 infection causes mild acute disease in the majority of those infected, but progresses to severe and fatal disease in a subset [1]. Extensive work has been done to elucidate the precise events responsible for disease progression in COVID-19. Two specific early events that accurately predict COVID-19 disease progression in humans have been independently identified. These events are (1) the early interaction between SARS-CoV-2 and megakaryocytes (MKs) in the bone marrow [2,3], and (2) the early production of a distinct form of non-neutralizing antibody against SARS-CoV-2 spike protein that drives the immunopathology of severe disease [4–6].

Multiple studies have shown that the quality of antibodies produced early on in response to SARS-CoV-2 spike protein differs between individuals, and the production of a distinct form of anti-spike IgG early in the illness accurately predicts disease progression [4–6]. In two prospective cohorts of patients with early COVID-19, the glycosylation pattern of the Fc component of the IgG formed against the spike protein was shown to determine the fate of these outpatients [4,6–8]. Individuals who, due to unknown factors, produced IgG antibodies with low core fucosylation against the spike protein (afucosylated anti-spike IgG) early in the illness as outpatients were at significantly higher risk to progress to hospitalization and severe COVID-19 in the ensuing days [4,6,8]. Those who instead formed anti-spike IgG antibodies with a normal level of core fucosylation (fucosylated anti-spike IgG) mounted an appropriate immune response to SARS-CoV-2 early on, experienced a mild disease course, and had a significantly lower risk of progression to severe COVID-19 [4–6].

Mechanistically, low core fucosylation of anti-spike IgG antibodies in COVID-19 is known to induce a potent pro-inflammatory state of cytokine production and lung immunopathology driven by intense FcγR-mediated stimulation of macrophages, monocytes, natural killer cells, and platelets [5,6]. Intriguingly, in individuals who produced afucosylated IgG against the spike protein (who later on progressed to severe disease), the low core fucosylation was limited to IgG produced against the spike protein which is a surface viral antigen and expressed on the surface membrane of infected host cells, whereas the degree of core fucosylation remained largely intact at a normal level for IgG formed against the viral nucleocapsid protein which lacks surface expression [4]. This is in alignment with instances of afucosylated IgG formation in other disorders, where afucosylated IgG production is seen in response to only surface antigens of enveloped viruses and to surface alloantigens of host blood cells such as platelets and RBCs [2,4,8]. Therefore, the capability of spike protein to be expressed on the surface of the viral envelope, and more importantly on the surface membrane of infected host cells, is hypothesized to play a key role and provide a specific antigen-presentation context that induces low core fucosylation of the humoral immune response to the spike protein in certain individuals [8]. However, spike protein is found abundantly expressed on the surface membrane of various host cells in the respiratory tract during SARS-CoV-2 infection, raising the question of whether specific cell populations expressing spike protein on their surface membrane may be required in order to trigger such aberrant afucosylated response that drives the immunopathology of severe COVID-19.

The body of knowledge in regards to afucosylated responses is currently limited. In the disorder of Fetal and Neonatal Allo-Immune Thrombocytopenia (FNAIT), afucosylated IgG are formed in reaction to alloantigens expressed on the platelet membrane and drive the immunopathology and thrombocytopenia in the fetus/newborn [9]. A hypothesis emanating from this phenomenon proposes that the expression of spike protein on the platelet membrane in COVID-19 may similarly induce formation of afucosylated anti-spike IgG antibodies and drive the immunopathology of severe illness. While it is known that whole and infectious SARS-CoV-2 is found within platelets in severe COVID-19, multiple studies utilizing electron microscopy and immunogold labeling of the spike protein have found the spike protein to be localized only in the cytoplasmic lumen of the infected platelets, and not expressed on the platelet membrane [3,4]. Bone marrow MKs, however, have been shown to be infected by SARS-CoV-2 as well, and are believed to be the most likely source of virus-infected platelets in COVID-19 [3,4]. In fact, virus-infected MKs were shown to transfer viral antigens including the spike protein to emerging platelets during thrombopoiesis [2]. Moreover, virus-infected MKs found early in the illness were predictive of disease progression to severe COVID-19 similar to the predictive nature of early afucosylated anti-spike IgG antibodies [2,3]. Thus far, however, no links between these two predictive factors have been discovered.

Based on the above lines of evidence, we hypothesized that, although the spike protein is not found expressed on the membrane of circulating infected platelets, spike protein expression on the membrane of platelet progenitors may occur during thrombopoiesis as infected MKs transfer viral antigens including the spike protein to budding pro-platelets. Surface expression of the spike protein on the membrane of pro-platelets during thrombopoiesis may then provide the specific antigen-presentation context that triggers the formation of afucosylated IgG against the spike protein, in a manner resembling the afucosylated response seen in reaction to platelet alloantigens implicated in the pathogenesis of FNAIT [9]. The production of afucosylated anti-spike IgG antibodies may then drive the immunopathology of severe COVID-19 by inducing a potent pro-inflammatory cascade via FcγR-mediated stimulation of macrophages, monocytes, natural killer cells, and platelets. This hypothesis was put to test in this work.

## Methods

After obtaining informed consent and with the approval of the local ethics committee (No. PR-1113–22), MKs were isolated from the bone marrow of a cohort of human subjects infected by SARS-CoV-2 who were experiencing severe COVID-19. The isolated MKs were transferred to an appropriate culture medium containing thrombopoietin and allowed to develop and differentiate under favorable growth conditions. The presence of SARS-CoV-2 was confirmed in the isolated MKs using real-time PCR. Isolated MKs confirmed to harbor SARS-CoV-2 were then injected intravenously into K18-hACE2 transgenic mice (RRID: IMSR_JAX:034860). Mice were monitored for signs of infection, disease, or behavioral changes. Blood samples were taken to assess the response to injection of the mice with virus-infected human MKs, and to quantify plasma levels of afucosylated anti-spike IgG antibodies and other molecules (serotonin and Platelet Factor 4 (PF4)) by ELISA. Once mice were euthanized, lung sections were prepared and processed with routine hematoxylin and eosin (H&E) staining and histopathological analysis was performed.

## Results

### Pro-platelets express SARS-CoV-2 spike protein on their surface membrane during maturation from virus-infected human MKs

MKs were isolated from the bone marrow of four human subjects infected by SARS-CoV-2 who were experiencing severe COVID-19. Mean duration from the onset of severe illness to bone marrow biopsy was 3.2±0.85 days. The isolated human MKs were confirmed to harbor SARS-CoV-2 using real-time PCR. Under favorable growth conditions in presence of thrombopoietin, virus-infected human MKs were observed to develop and differentiate to pro-platelets. Pro-platelets derived from virus-infected human MKs expressed SARS-CoV-2 spike protein on their surface membrane during their maturation. As shown in Figure 1, the periphery of pro-platelets derived from virus-infected human MKs is marked in green with anti-α-tubulin, while SARS-CoV-2 spike protein expression is marked in red. As observed, spike protein is expressed on the surface membrane of maturing pro-platelets emerging from virus-infected human MKs. Visualization was performed by confocal microscopy (teaser) after pro-platelet fixation.

**Figure 1.**
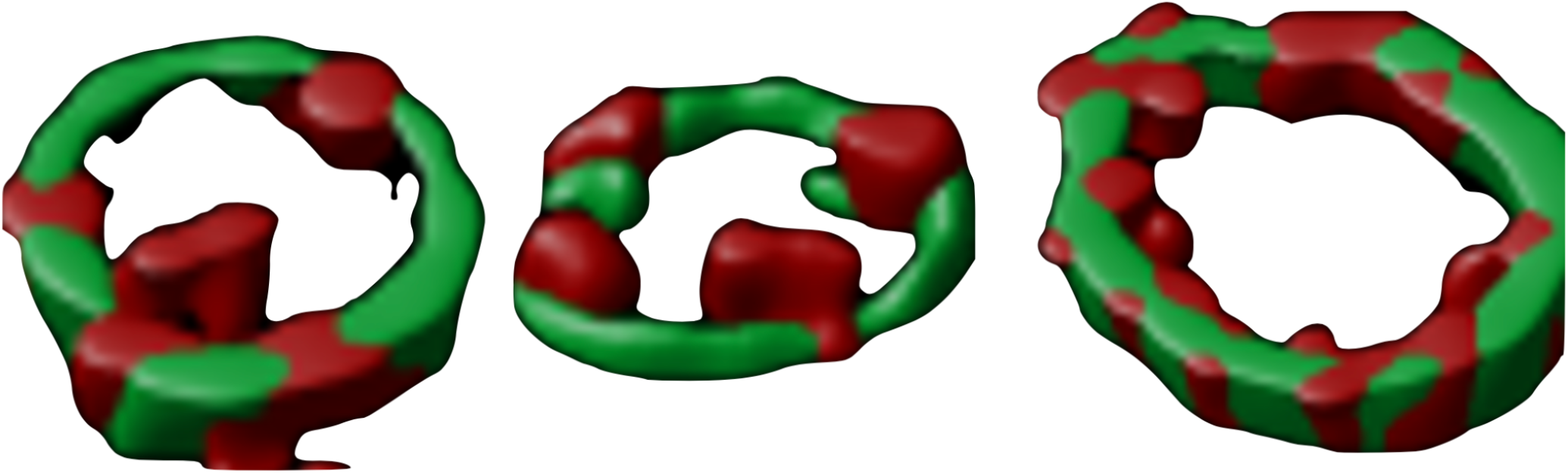
In green: α-tubulin marks the periphery of pro-platelets emerging from MKs. In red: SARS-CoV-2 spike protein localized on the surface of pro-platelets during maturation.

### Afucosylated anti-spike IgG production occurs in response to intravenous injection of SARS-CoV-2-infected human MKs, and promotes platelet activation, lung injury, and death in mice

Isolated MKs from human bone marrow confirmed to harbor SARS-CoV-2 were injected intravenously into K18-hACE2 transgenic mice (RRID: IMSR_JAX:034860). Injection of virus-infected human MKs into the transgenic mice significantly promoted the formation afucosylated IgG antibodies against the spike protein (Figure 2). The mortality rate in the infected mice was proportional to the level of afucosylated anti-spike IgG produced (Table 1). This finding suggests that the afucosylated anti-spike IgG antibodies were pathogenic and contributed significantly to the severity of COVID-19 in mice (as observed in humans [4,6,8]).

**Table 1.** The mortality rate in mice infected with SARS-CoV-2-infected human MKs compared to non-infected mice. Unpaired Student *t* test was used to calculate p-values. Bold numbers indicate statistical significance at p-value ≤0.05.

|  | Infected mice | Non-infected mice | p-value |
| --- | --- | --- | --- |
| Mortality rate after 3 days (%) | 33.3 | 8.3 | **0.02** |
| Mortality rate after 7 days (%) | 58.3 | 8.3 | **0.01** |

**Figure 2.**
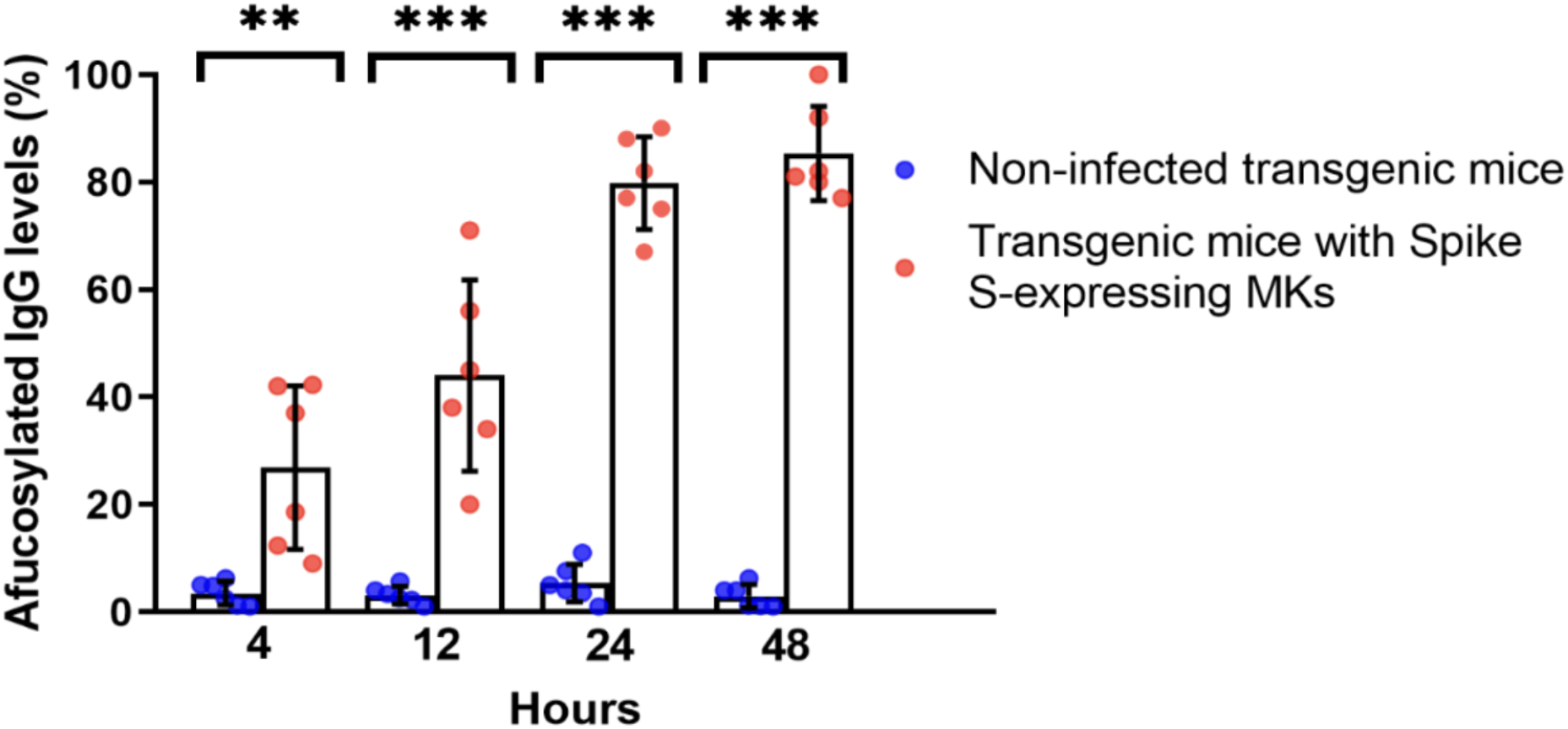
K18-hACE2 transgenic mice infected with SARS-CoV-2-infected human MKs produced significantly elevated levels of afucosylated anti-spike IgG antibodies.

The plasma levels of PF4 and serotonin (Figure 3), as surrogate soluble markers of platelet activation, were significantly higher in the infected mice (marked in red) compared with the lower levels of PF4 and serotonin in the uninfected mice (marked in blue).

**Figure 3.**
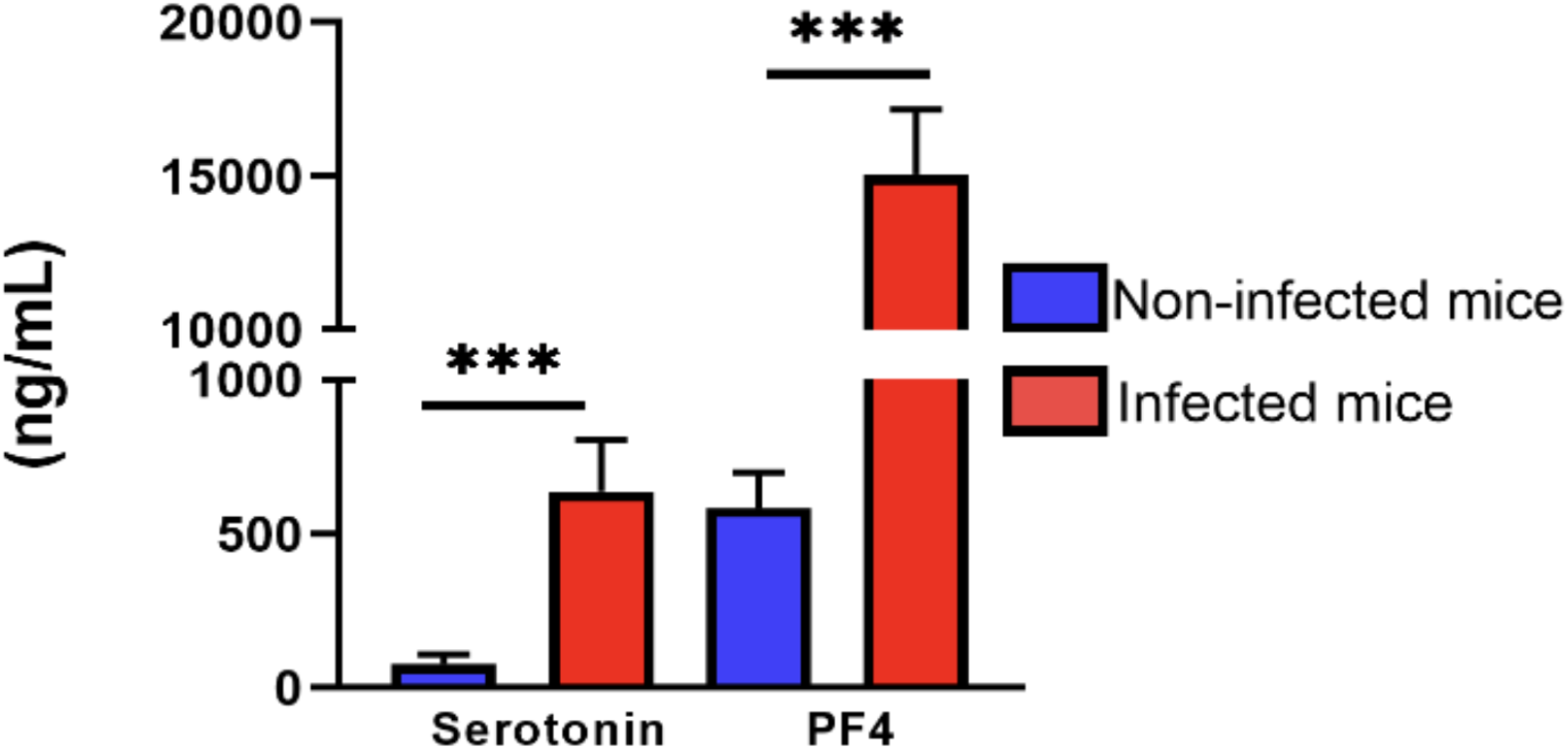
The plasma levels of serotonin and PF4 are significantly elevated in mice infected with SARS-CoV-2-infected human MKs compared to non-infected mice. Paired Student *t* test was used to calculate p-values. ^***^ indicates statistical significance at p-value ≤ 0.05.

Histopathological analysis of the lung tissue after H&E staining was carried out on euthanized mice as shown in Figure 4. Panel A shows a control with normal lung tissue. Panels B and C are cross-sections 4 hours and 24 hours post-infection of the transgenic mice with virus-infected human MKs, demonstrating inflammatory cell infiltrate, thrombus formation in the pulmonary vascular lumen, and distortion of the lung tissue.

**Figure 4.**
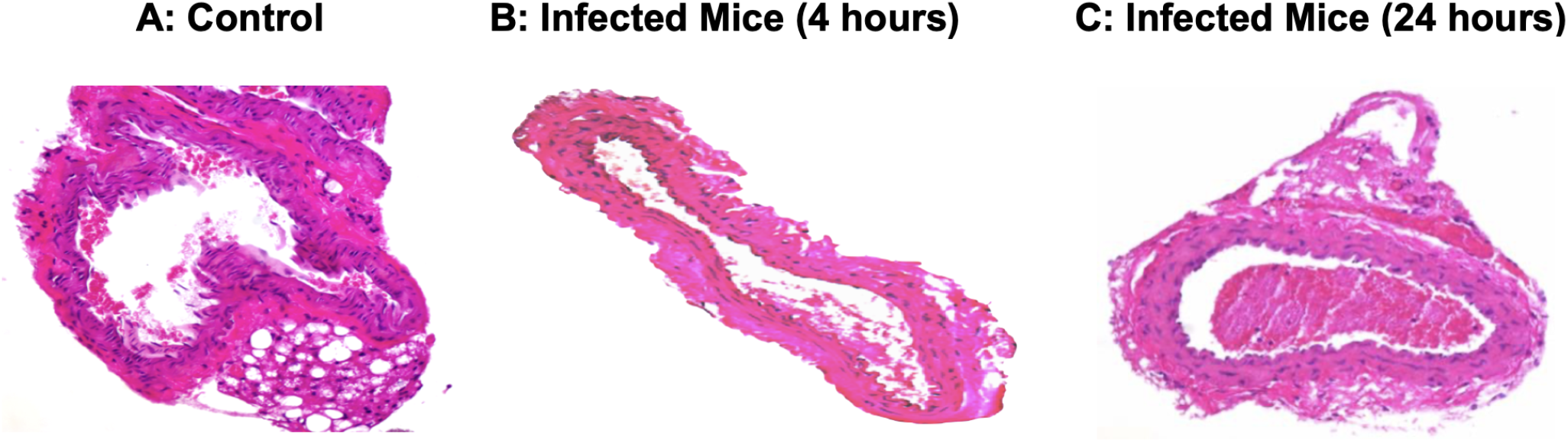
Infection of transgenic mice with SARS-CoV-2-infected human MKs induces acute lung injury and the formation of pulmonary vascular thrombosis.

## Discussion

This investigation adds to the existing body of knowledge that human MKs residing in the bone marrow become infected by SARS-CoV-2 in severe and fatal COVID-19. All prior studies to date demonstrating clear evidence of MK infection by SARS-CoV-2 were performed on autopsy tissues from deceased COVID-19 donors [2,3,11,12]. Due to the postmortem nature of these studies, no specific assessment could be made regarding the timing of bone marrow MK infection, or whether bone marrow MK infection by SARS-CoV-2 was an early event or, vice versa, a byproduct of the multiorgan failure in the fatal stage of the illness. To the best of our knowledge, this is the first study that demonstrates direct evidence of bone marrow MKs exhibiting infection by SARS-CoV-2 in living donors relatively early in the course of severe illness (3.2±0.85 days from the onset of severe illness). This finding is in agreement with the study by Xhu *et al*. which provided indirect evidence of virus-infected bone marrow MKs early in the illness, a finding that also accurately predicted COVID-19 disease progression and mortality in a human cohort [3].

Additionally, we were able to demonstrate that during differentiation of virus-infected human MKs, SARS-CoV-2 spike protein is expressed on the surface of the emerging pro-platelets’ membrane. Although virus-infected MKs were previously shown to transfer viral antigens including the spike protein to emerging platelets during thrombopoiesis [2], prior studies did not find specific evidence of surface expression of spike protein on the membrane of virus-infected platelets, MKs, or other platelet progenitors [3,4]. The finding of spike protein expression on the membrane of emerging pro-platelets in this work is of potentially high relevance to the well-described afucosylated IgG immunopathology responsible for COVID-19 disease progression [4–6].

Larsen *et al*. have demonstrated that afucosylated IgG responses are not seen in response to soluble proteins in plasma, internal proteins of enveloped viruses (including the nucleocapsid protein of SARS-CoV-2), or proteins of non-enveloped viruses [4]. The prevailing hypothesis is that afucosylated IgG responses are triggered by foreign antigens expressed on the surface membrane of host blood cells, either when these host blood cells become infected by enveloped viruses, or in response to platelet (or RBC) alloantigens as it occurs in FNAIT. The antigen-presenting context that leads to such altered glyco-programming and distinctly afucosylated IgG antibodies, however, has never been demonstrated in live animals [4,8].

For the first time to the best of our knowledge, we have demonstrated that exposure in the bloodstream to human bone marrow MKs infected by SARS-CoV-2 promotes the formation of afucosylated IgG antibodies against the spike protein in mice. In a manner immunologically resembling afucosylated responses to platelet membrane alloantigens in FNAIT, spike protein expression on the membrane of pro-platelets, as demonstrated in this work, may induce a similar antigen-presenting context that leads to altered glyco-programming of the humoral response and results in the formation of afucosylated anti-spike IgG antibodies. Lastly, we have demonstrated that afucosylated anti-spike IgG production, as induced by bloodstream exposure to virus-infected human MKs, promotes soluble markers of platelet activation, induces pulmonary vascular thrombosis and acute lung injury, and is associated a fatal disease course in mice resembling that of severe COVID-19 in humans.

In summary, this work provides a plausible mechanism that ties together the infection of bone marrow MKs by SARS-CoV-2 as a likely trigger for the early production of non-neutralizing and pathogenic afucosylated anti-spike IgG antibodies. The infection of bone marrow MKs [2,3] and the formation of afucosylated anti-spike IgG antibodies [4–6] are demonstrated to precede and drive COVID-19 disease progression in humans. Based on our observations, the intravenous injection of virus-infected human MKs alone, without inhalation of SARS-CoV-2 virus, was sufficient to significantly promote the formation of such afucosylated anti-spike IgG antibodies and cause pulmonary vascular thrombosis, acute lung injury and death in the mice.

A similar sequence of events may be at play in human cases of severe COVID-19, whereby a breach of early innate and adaptive immune defenses by SARS-CoV-2 introduced via the respiratory tract may allow dissemination of the virus to the bone marrow niche in susceptible individuals. Once it has reached the bone marrow, SARS-CoV-2 infects the bone marrow MKs as demonstrated by five recent studies including the most direct human evidence presented in this work [2,3,11,12]. Virus-infected bone marrow MKs may subsequently undergo differentiation to infected pro-platelets that express spike protein on their membrane, culminating (as demonstrated in mice in this work) in the production of non-neutralizing, pathogenic afucosylated anti-spike IgG antibodies prior to the development of an appropriate humoral response consisting of neutralizing, fucosylated anti-spike IgG antibodies that are known to take longer time to form in human cases that progress to severe COVID-19 [4].

The novel findings and the proposed mechanisms in this work raise several lines of inquiry that will be the focus of our future work, including most urgently (1) a deeper investigation to elucidate the interaction between surface-expressed viral antigens of various enveloped viruses and human MKs, and (2) the signaling pathways that trigger altered glyco-programming in the humoral immune effector cells such as B cells that promote low core fucosylation of IgG antibodies against viral antigens presented on the membrane of platelet progenitors. This understanding would likely facilitate the identification of therapeutic interventions to prevent the formation of such pathogenic afucosylated antibodies, which not only play a significant role in the morbidity and mortality associated with COVID-19 and dengue fever [10], but also have described immunologic roles in HIV, malaria, as well as in the design of potent cancer therapeutics [8].

## Data Availability Statement

The original contributions presented in the study are included in the article, and further inquiries can be directed to the corresponding author.

## Ethical Approval Statement

Written informed consent was obtained from the individuals for the publication of any data included in this article.

## Authors’ Contribution

F. J., F. G. and Y. Z. designed the research. M. M., N. Z., A. N., F. G. and Y. Z. performed the research and statistical analyses. F. J., L. K., F. G., and Y. Z. analyzed the data. F. J., M. K., F. G. and Y. Z. contributed to discussion and revised the manuscript. F. J., M. M., F. G. and Y. Z. wrote the manuscript. All authors read and approved the paper.

## References

[1] Bouadma L, Lescure F-X, Lucet J-C, Yazdanpanah Y, Timsit J-F. Severe SARS-CoV-2 infections: practical considerations and management strategy for intensivists. Intensive Care Med 2020;46:579–82. https://doi.org/10.1007/s00134-020-05967-x.

[2] Fortmann SD, Patton M, Frey BF, Tipper JL, Reddy SB, Vieira CP, et al. Circulating SARS-CoV-2+ Megakaryocytes Associate with Severe Viral Infection in COVID-19. Blood Adv 2023:bloodadvances.2022009022. https://doi.org/10.1182/bloodadvances.2022009022.

[3] Zhu A, Real F, Capron C, Rosenberg AR, Silvin A, Dunsmore G, et al. Infection of lung megakaryocytes and platelets by SARS-CoV-2 anticipate fatal COVID-19. Cell Mol Life Sci CMLS 2022;79:365. https://doi.org/10.1007/s00018-022-04318-x.

[4] Larsen MD, De Graaf EL, Sonneveld ME, Plomp HR, Nouta J, Hoepel W, et al. Afucosylated IgG characterizes enveloped viral responses and correlates with COVID-19 severity. Science 2021;371:eabc8378. https://doi.org/10.1126/science.abc8378.

[5] Hoepel W, Chen H-J, Geyer CE, Allahverdiyeva S, Manz XD, De Taeye SW, et al. High titers and low fucosylation of early human anti–SARS-CoV-2 IgG promote inflammation by alveolar macrophages. Sci Transl Med 2021;13:eabf8654. https://doi.org/10.1126/scitranslmed.abf8654.

[6] Chakraborty S, Gonzalez JC, Sievers BL, Mallajosyula V, Chakraborty S, Dubey M, et al. Early non-neutralizing, afucosylated antibody responses are associated with COVID-19 severity. Sci Transl Med 2022;14:eabm7853. https://doi.org/10.1126/scitranslmed.abm7853.

[7] Chakraborty S, Gonzalez J, Edwards K, Mallajosyula V, Buzzanco AS, Sherwood R, et al. Proinflammatory IgG Fc structures in patients with severe COVID-19. Nat Immunol 2021;22:67–73. https://doi.org/10.1038/s41590-020-00828-7.

[8] Oosterhoff JJ, Larsen MD, Van Der Schoot CE, Vidarsson G. Afucosylated IgG responses in humans – structural clues to the regulation of humoral immunity. Trends Immunol 2022;43:800–14. https://doi.org/10.1016/j.it.2022.08.001.

[9] Kapur R, Kustiawan I, Vestrheim A, Koeleman CAM, Visser R, Einarsdottir HK, et al. A prominent lack of IgG1-Fc fucosylation of platelet alloantibodies in pregnancy. Blood 2014;123:471–80. https://doi.org/10.1182/blood-2013-09-527978.

[10] Bournazos S, Vo HTM, Duong V, Auerswald H, Ly S, Sakuntabhai A, et al. Antibody fucosylation predicts disease severity in secondary dengue infection. Science. 2021 Jun 4;372(6546):1102–1105. DOI: 10.1126/science.abc7303. PMID: 34083490; PMCID: PMC8262508.

[11] Barrett TJ, Bilaloglu S, Cornwell M, Burgess HM, Virginio VW, Drenkova K, et al. Platelets contribute to disease severity in COVID-19. J Thromb Haemost. 2021 Dec;19(12):3139–3153. DOI: 10.1111/jth.15534. Epub 2021 Sep 29. PMID: 34538015; PMCID: PMC8646651.

[12] Gray-Rodriguez S, Jensen MP, Otero-Jimenez M, Hanley B, Swann OC, Ward PA, et al. Multisystem screening reveals SARS-CoV-2 in neurons of the myenteric plexus and in megakaryocytes. J Pathol. 2022 Jun;257(2):198–217. DOI: 10.1002/path.5878. Epub 2022 Mar 31. PMID: 35107828; PMCID: PMC9325073.

